# Fast Interneuron Dysfunction in Laminar Neural Mass Model Reproduces Alzheimer’s Oscillatory Biomarkers

**DOI:** 10.1101/2025.03.26.645407

**Authors:** Roser Sanchez-Todo, Borja Mercadal, Edmundo Lopez-Sola, Maria Guasch-Morgades, Gustavo Deco, Giulio Ruffini

**Affiliations:** Brain Modeling Department, Neuroelectrics Barcelona, 08035 Barcelona, Spain; Center of Brain and Cognition, Computational Neuroscience Group, Department of Information and Communication Technologies, Universitat Pompeu Fabra, Barcelona, Spain

## Abstract

Alzheimer’s disease (AD) is characterized by a progressive cognitive decline underpinned by disruptions in neural circuit dynamics. Early-stage AD is associated with cortical hyperexcitability, whereas later stages exhibit oscillatory slowing and hypoactivity, a progression observable in electrophysiological spectral characteristics. While previous studies have linked these changes to the dysfunction of fast-spiking parvalbumin-positive (*PV*) interneurons and neuronal loss associated with amyloid-beta (Aβ) and hyperphosphorylated tau (hp-*τ*) pathology, the precise mechanistic relationship between cellular and altered electrophysiology remains unclear. To study this relationship, we employed a Laminar Neural Mass Model that integrates excitatory and inhibitory neural populations within a biophysically informed columnar framework. The connectivity constant from *PV* cells to pyramidal neurons was gradually reduced to simulate the progressive neurotoxic effects of Aβ oligomers. Other model parameters were systematically varied to compare with existing modeling literature and also to simulate the effects of hp-*τ*. All model predictions were compared to empirical M/EEG findings in the literature. Our simulations of *PV* interneuron dysfunction successfully reproduced the biphasic electrophysiological progression observed in AD: an early phase of hyperexcitability with increased gamma and alpha power, followed by oscillatory slowing and reduced spectral power. Alternative mechanisms and model parameters, such as increased excitatory drive, failed to replicate the observed biomarker trajectory. Additionally, to reconcile the hypoactivity and decreased firing rates observed in advanced AD stages, we combined the *PV* dysfunction model with a disruption of the pyramidal cell populations that reflects the neurotoxicity induced by hp-*τ*. Although this additional mechanism is not necessary to reproduce oscillatory changes in the isolated neural mass, it is crucial for aligning the model with evidence of reduced firing rates, metabolic activity, and cell loss and will enhance its applicability in future whole-brain modeling studies. These results support the hypothesis that at the local level, *PV* interneuron dysfunction is a primary driver of cortical electrophysiological alterations, while pyramidal neuron loss underlies later-stage severe hypoactivity. Our model provides a mechanistic framework for interpreting excitation-inhibition imbalance across AD progression, demonstrating the value of biophysically constrained models for interpreting electrophysiological biomarkers.

**Author summary:** Alzheimer’s disease (AD) is not just a disorder of memory—it is a disease of brain networks. Before widespread neuronal loss occurs, the brain enters a state of hyperexcitability, which may contribute to disease progression. This hyperactivity is thought to arise from the selective dysfunction of inhibitory interneurons, particularly parvalbumin-positive (*PV*) interneurons, which play a crucial role in maintaining balanced brain activity. However, as the disease advances, a dramatic shift occurs, with neurons becoming progressively less active, leading to network breakdown. Understanding how this transition unfolds is essential for identifying new targets for early intervention.

In this study, we developed a computational model of AD that represents the effects of amyloid-beta (Aβ) oligomers and hyperphosphorylated tau (hp-*τ*) on neural circuits in a biologically meaningful way. Our model reproduces key features of M/EEG biomarkers, demonstrating that *PV* interneuron dysfunction leads to early hyperexcitability and the electrophysiological oscillatory changes characteristic of AD, while pyramidal cell pathology is necessary to drive later hypoactivity and network failure.

By bridging molecular pathology mechanisms with mesoscale neural activity, our model provides a powerful tool for studying AD-related circuit dysfunction. It also highlights the importance of *PV* interneurons as a potential therapeutic target, paving the way for biologically informed interventions that could alter the course of the disease.

## Introduction

Millions of people worldwide are affected by Alzheimer’s disease (AD), the most common type of dementia. It is characterized by the presence of extracellular plaques of amyloid beta (Aβ) proteins and intracellular fibrils of hyperphosphorylated tau protein (hp-*τ*). Clinically, AD is defined by a gradual decline in cognitive function, progressing along a continuum from an asymptomatic preclinical stage to advanced dementia.^1, 2^ As the global population ages, the impact of AD on society and economics is intensifying, emphasizing the urgency of finding early solutions to slow disease progression and improve patient outcomes.^2, 3^ Identifying the brain changes across the different stages of the disease is essential for enabling early diagnosis and guiding targeted treatments.^4^

The criteria for diagnosing AD in its various stages has been revised during the past decade by the National Institute on Aging and the Alzheimer’s Association (NIA-AA).^5–11^ In their latest update,^2^ they proposed a framework combining biological and clinical stages, aligning with other well-known classifications.^1^ This framework divides the disease continuum into three primary stages: preclinical, Mild Cognitive Impairment due to AD (MCI, also known as prodromal AD), and AD dementia. The preclinical stage is further divided into three substages. Stage 0 includes individuals with no cognitive symptoms but genetic predisposition. Stage 1 features Aβ deposition without cognitive decline. Stage 2 involves subtle cognitive decline, still within normal limits, often accompanied by hp-*τ* deposition in the medial and temporal regions. At this point, individuals may experience Subjective Cognitive Decline (SCD)—self-reported worsening of cognitive function despite normal performance on standard neuropsychological tests.^12^ SCD is considered a very early marker of risk, associated with an increased likelihood of progression to MCI and Alzheimer’s dementia. In MCI, there is objective evidence of cognitive impairment, with an early functional impact on daily life. This stage often correlates with an increased burden of Aβ and hp-*τ* pathology in cortical areas, signaling a transition toward AD dementia. Finally, the AD stage, diagnosed in its mild, moderate, and severe forms, is marked by progressively worsening cognitive and functional impairment and increased deposition of Aβ and hp-*τ* . See Fig. 1a for an overview of the progression of the main biomarkers throughout the disease stages.

**Fig 1.**
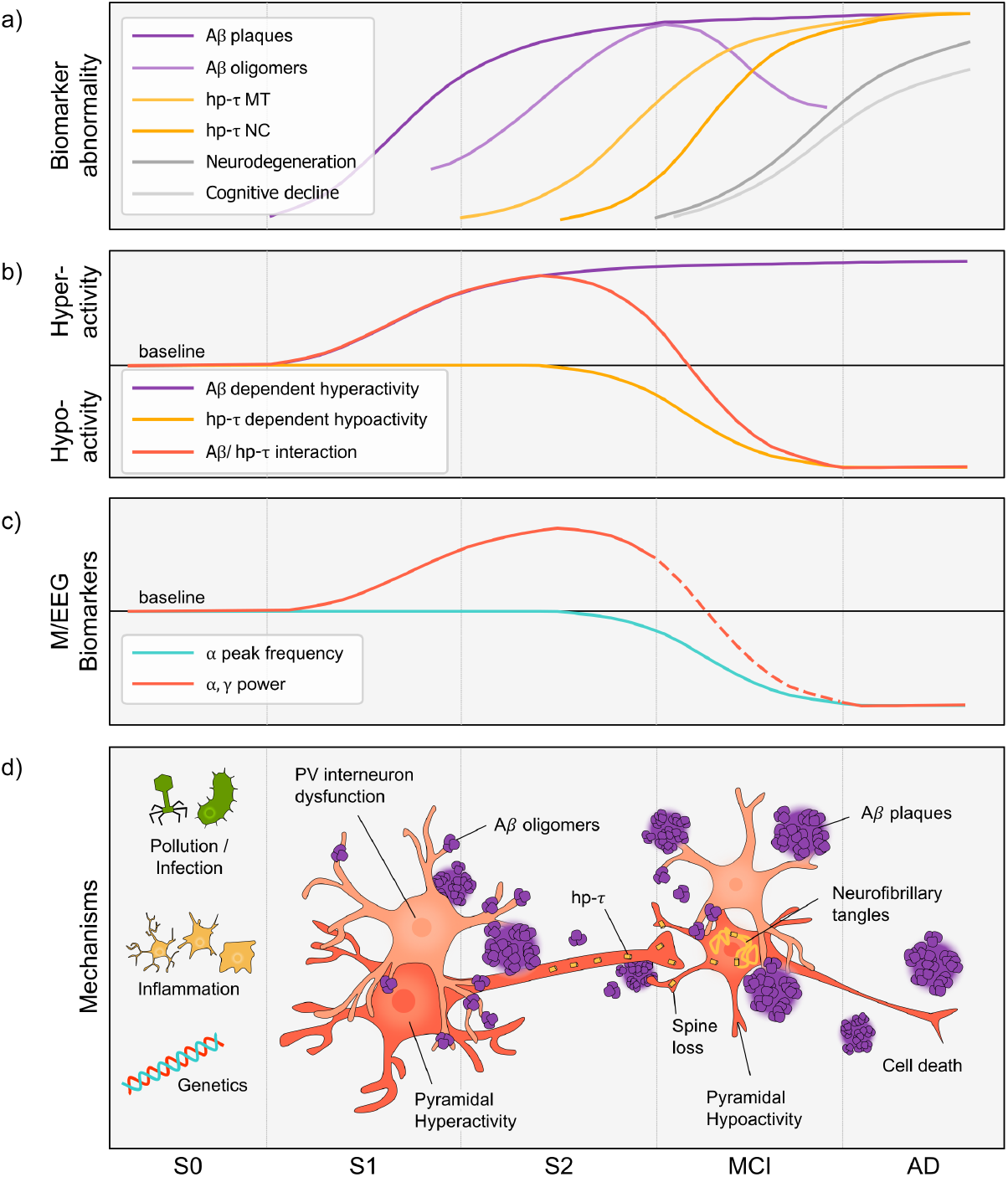
a) Biomarkers abnormality throughout the different stages of AD. Modified from Jack et al. (2024)^2^ and Blomeke et al. (2024).^37^ The deposition of Aβ oligomers may follow distinct trajectories depending on genetic factors.^37^ MT refers to medial temporal lobe, and NC neocortical. b) Model of Aβ and hp-*τ* effect on the hypo- and hyper-activity and their interaction (adapted from Harris et al. 2020^40^). c) M/EEG Biomarkers progression, specifically alpha and gamma biomarkers, as inferred from the literature. d) Overview of the main mechanisms throughout the disease progression. The boundaries between clinical stages are to be seen as an approximation to account for the variance in diagnostic methods across papers.

Some hypotheses suggest that AD may originate from viral,^13–18^ bacterial,^19–22^ or fungal^23^ infection, or environmental insults like air pollution,^24–31^ influenced by genetic risk factors such as the presence of apolipoprotein or the APOE4 allele.^32^ This immune response elevates the levels of soluble Aβ oligomers as antimicrobial agents,^22, 33–36^ which peak during early stages of AD^37^ and exert neurotoxic effects, disrupting cortical circuit excitation/inhibition homeostasis.^38–42^ This process initiates a vicious cycle, as increased neural activity, in turn, fosters additional Aβ oligomer production.^38, 43^ The oligomer form of Aβ prior to plaque formation is thought to be particularly toxic for Parvalbumin positive (*PV*) interneurons,^44–48^ which are critical for driving gamma oscillations linked to cognitive functions.^44, 49^ Aβ oligomers consolidate then into plaques, subsequently accelerating hp-*τ* spreading,^50^ particularly when mutant forms of hp-*τ* are present.^51^ hp-*τ*, in turn, seems to target excitatory neurons, specifically pyramidal cells.^52–55^ The progressive accumulation of Aβ and hp-*τ* exacerbates synaptic dysfunction, resulting in spine loss, neuronal death, and widespread hypoactivation in later stages of the disease.^40, 47, 50^ See Fig. 1d for an overview of the main mechanisms over the disease stages, which map to biomarkers and neutral activity shown in Fig. 1a-c.

A range of biomarkers are used in research and clinical practice for AD, including the analysis of cerebrospinal fluid (CSF) to detect levels of Aβ and hp-*τ*, amyloid and Fluorodeoxyglucose Positron Emission Tomography (FDG-PET) scans, and Magnetic Resonance Imaging (MRI) volumetry.^2, 56^ Early-stage biomarkers are crucial because biological changes, such as Aβ deposition, can begin years before clinical symptoms appear. Electroencephalography (EEG) and Magnetoencephalography (MEG) are non-invasive, cost-effective, and repeatable methods for monitoring disease progression, and are good but underutilized candidates to derive early-stage biomarkers. Unlike more expensive or invasive techniques like PET or CSF analysis, M/EEG can detect early synaptic dysfunction due to their high temporal resolution for measuring neural activity and brain oscillations.^2, 56–59^ In what follows, we will explain the main findings regarding alpha and gamma frequencies throughout the disease progression, which are shown in Fig. 1c and in Table 1.

**Table 1.** Electrophysiological changes observed through the Alzheimer’s disease progression.

| Feature | Progression across stages |
| --- | --- |
| Alpha Peak Frequency | • Preclinical (S0, S1, S2): No consistent evidence of alterations compared to healthy controls.<br>• MCI: Slowing of the alpha peak frequency is reported.<sup>60,61,65,67</sup><br>• AD: Pronounced slowing in line with widespread neural atrophy and disease severity.<sup>73–75</sup> |
| Alpha Band Power | • Preclinical: Possible increases,<sup>4,60</sup> though some studies indicate reductions in S2.<sup>61</sup><br>• MCI: Conflicting evidence, both increases<sup>60</sup> and decreases.<sup>65,66</sup><br>• AD: Progressive decreases in alpha power become prominent.<sup>68–70,73–76</sup> |
| Gamma Band Power | • Preclinical: One report of increases in power.<sup>4</sup><br>• MCI: Findings differ, with evidence of both increases<sup>70</sup> and decreases.<sup>69</sup><br>• AD: A generalized reduction of gamma power.<sup>68,69,77</sup> |

In the preclinical stages of Alzheimer’s disease (S0, S1, and S2), evidence has highlighted neural hyperexcitability in humans, as observed through M/EEG biomarkers (Fig. 1b). Gaubert et al. (2019)^4^ reported increased gamma power in cognitively normal individuals with Aβ deposition. This was accompanied by heightened functional connectivity (FC) in the alpha band, indicative of hyperactivity within these frequency ranges. Similarly, Nakamura et al. (2018)^60^ observed elevated alpha power in cognitively normal subjects with Aβ deposition. However, some findings suggest a contrasting reduction in alpha band activity.^61^ This simultaneous increase in alpha-band spectral power across multiple brain regions is thought to reflect an imbalance between excitation and inhibition, contributing to neural hyperactivity and network disruptions.^56, 62^ Such disruptions are correlated with pathological biomarkers, including plasma hp-*τ*, and are linked to cognitive decline.^63, 64^ These findings support the hypothesis that early-stage Alzheimer’s disease is characterized by disruptions in excitation-inhibition balance.

In the prodromal or MCI due to AD stage, there is mixed evidence regarding excitation-inhibition imbalance, with some studies reporting hyperexcitability and increased functional coupling or elevated spectral power in low-frequency bands, while others report opposite findings, perhaps reflecting the difficulty in differentiating preclinical and MCI states of AD. For the alpha band, most studies report reduced power compared to healthy controls,^61, 65, 66^ though some studies have observed increased power.^60^ There is consistent evidence of spectral slowing, reflected in a lower alpha peak frequency.^60, 61, 65, 67^ Findings on high-frequency activity in Alzheimer’s disease remain inconclusive, with some studies reporting reductions and slowing,^68, 69^ while others observe increases in power,^70^ highlighting conflicting evidence on whether these oscillations are suppressed or enhanced. In terms of network disruptions, early stages are characterized by anterior alpha-band coherence and reduced posterior functional connectivity.^64, 71^ Notably, abnormally increased alpha-band coherence has been shown to predict progression from MCI to AD and is linked to excitation-inhibition imbalance-driven circuit disruptions.^72^ These findings highlight the complexity of neural activity alterations in the prodromal stages of AD. Despite conflicting findings in higher frequencies, consistent evidence of spectral slowing and network disruptions underscores the potential of M/EEG biomarkers for identifying early AD and tracking its progression.

The later stages of Alzheimer’s disease (mild, moderate, and severe) are marked by extensive neuronal loss and progressive neurodegeneration. Reductions in both alpha^61, 66, 68, 70, 73–76^ and gamma power^68, 69, 77^ are accompanied by increases in slower rhythms, particularly delta and theta.^66, 68, 70, 74, 75^ These spectral shifts align with widespread cortical atrophy, excitation-inhibition imbalance, and network synchronization disruptions, contributing to the profound cognitive and functional decline characteristic of advanced AD. These oscillatory changes in alpha and gamma power, alongside the rise in slower rhythms, highlight the progressive neural disruptions in AD and underscore the utility of EEG biomarkers in capturing the disease’s impact on cortical dynamics and cognitive function.

In order to map pathology to our modeling framework, we first review the current understanding of AD etiology. Neurophysiological evidence suggests that Aβ oligomers have an initial effect, which is the increase in the activity of neurons^40, 78–80^ (Fig. 1b). Among the proposed mechanisms underlying this hyperexcitability, our primary hypothesis is that reduced inhibitory activity plays a central role, especially diminished fast GABAergic signaling. However, other contributing factors, including increased excitatory drive and reactive oxygen species (ROS) toxicity, have also been suggested as key mechanisms.^40, 78^ In the later stages of the disease, hypoactivity is induced by hp-*τ* deposition and cell death (Fig. 1b).

Our primary mechanistic hypothesis is that the hyperexcitability observed in animal models of AD arises primarily from impaired inhibition and is closely linked to soluble Aβ oligomers rather than plaques. Busche et al. 2008^81^ showed that hyperactive neurons, particularly those near plaques, exhibit increased calcium transients, suggesting reduced inhibitory control. Diazepam, a GABAa receptor modulator, restored inhibition, implicating defective synaptic inhibition as the primary driver of hyperactivity rather than intracellular calcium release. Ren et al. 2018^82^ confirmed that Aβ oligomers depress inhibitory transmission, particularly in fast-spiking interneurons, exacerbating excitability. Moreover, Busche and Konnerth 2015^43, 80^ concluded that hyperactivity is an early dysfunction in AD caused by soluble Aβ oligomers, with plaques being nonessential. *PV* interneurons are disproportionately affected in AD. Garcia-Marin et al. 2009^47^ observed a loss of GABAergic perisomatic synapses on neurons near plaques (e.g., from *PV* cells), indicating a local reduction of inhibitory input. Verret et al. 2012^83^ and Martinez-Losa et al. 2018^48^ demonstrated that restoring impaired Nav1.1 channels in *PV* interneurons improved cognitive performance and restored gamma oscillations, a marker of network stability and cognition.^44^ Similarly, Chung et al. 2020^45^ showed that optogenetic activation of *PV* neurons selectively restored gamma power and resynchronized CA1 pyramidal cell spikes in Aβ oligomer-injected mice. Zhang et al. 2024^84^ reviewed evidence that loss of *PV* interneurons contributes to increased excitability and cognitive decline in AD. In summary, these findings support the idea that the electrophysiological changes across stages in AD are primarily driven by reduced inhibition by damaged interneurons, which results from the deleterious effects of soluble Aβ oligomers rather than plaques. Among inhibitory neurons, *PV* interneurons are the most affected, as they are particularly vulnerable to Aβ oligomers and the main responsible for gamma oscillations, which are altered in AD.^44^

Another mechanism that could contribute to hyperactivity and consequent cell damage generated by Aβ oligomers is the disruption of glutamate homeostasis and enhancement of excitatory neurotransmission. This can occur when astrocytes are impaired in their ability to reuptake glutamate, resulting in the accumulation of extracellular glutamate.^38^In vivo studies have demonstrated that Aβ oligomers reduce the function of excitatory amino acid transporter 2 (EAAT2/GLT-1), leading to extracellular glutamate accumulation and excessive excitatory drive.^38, 85, 86^ This reuptake impairment was found to be associated with lower EAAT2 levels and diffusion obstruction within astrocytic membranes, particularly in the CA1 region of the hippocampus, a site of early AD pathology.^38, 85, 87^ The consequent glutamate buildup enhances activation of NMDA (N-Methyl-D-Aspartate) receptors, leading to calcium overload and neuronal hyperactivity, which is exacerbated by the activity-dependent production of Aβ, forming a positive feedback loop that sustains excitotoxicity.^38, 88^ Additionally, Aβ oligomers can promote hyperexcitability by enhancing presynaptic glutamate release. Studies have shown that Aβ oligomers increase the probability of synaptic vesicle exocytosis by disrupting the synaptophysin/VAMP2 complex, thereby amplifying glutamate release at excitatory synapses.^89, 90^ Even small elevations in endogenous Aβ peptides have been found to accelerate vesicle exocytosis and increase the release probability of active neurons in hippocampal cultures.^91^ Furthermore, Aβ oligomers induced presynaptic calcium influx via Amyloid Precursor Protein (APP) homodimerization has been shown to enhance glutamate release, contributing to an overall increase in synaptic excitation.^92^ This excess glutamate can lead to “spillover,” activating extrasynaptic GluN2B-containing NMDA receptors, which have been implicated in neurotoxicity and synaptic dysfunction in AD.^93^ Notably, studies indicate that hyperactivity itself can promote further Aβ release, perpetuating a cycle of excitatory dysregulation.^94–96^

Neuronal hyperexcitability places significant metabolic strain on neurons, leading to increased production of ROS. Overactive neurons have elevated energy and oxygen demands to support excessive firing. This heightened metabolic demand overburdens mitochondria, the primary source of ROS in cells.^78^ Excess Ca^2+^ entering neurons during hyperexcitation is taken up by mitochondria, impairing their electron transport chain and stimulating ROS generation. Using neuronal cultures and entorhinal–hippocampal slices, researchers found that Aβ oligomers disrupted Ca^2+^ homeostasis and induced neuronal death via NMDA and AMPA (*α*-Amino-3-hydroxy-5-methyl-4-isoxazolepropionic) receptor activation, partially mediated by ROS generated from mitochondrial sources.^97, 98^

Together, these mechanisms illustrate how Aβ oligomers-driven disruption of glutamate regulation fosters persistent hyperexcitability, exacerbating neuronal stress and promoting disease progression. By increasing glutamate release and impairing its clearance, Aβ oligomers contribute to a pathological state of sustained excitatory drive, reinforcing the role of excitatory dysfunction as a crucial early feature of AD.

As AD progresses, hp-*τ* pathology becomes more prominent, damaging pyramidal neurons.^52, 55^ Emerging evidence suggests that hp-*τ* leads to neuronal hypoactivity, spine loss, and, ultimately, cell death. In vivo studies using transgenic mouse models overexpressing hp-*τ* have consistently shown that soluble hp-*τ* species are critical drivers of neuronal suppression, even in the absence of neurofibrillary tangles.^99, 100^ hp-*τ* disrupts neuronal firing patterns, reduces functional connectivity,^101^ and impairs synaptic function through mislocalization of hp-*τ* in dendritic spines, disrupting glutamate receptor trafficking^102^ and presynaptic vesicle release.^103^ Additionally, hp-*τ* induces significant downregulation of neuronal and synaptic genes, particularly those involved in glutamate signaling, further contributing to neuronal hypoactivity and spine loss.^55, 104–106^ Collectively, these hp-*τ* -induced deficits culminate in neuronal dysfunction and progressive neurodegeneration. Concomitantly, regional atrophy begins in medial temporal structures such as the hippocampus and extends to lateral temporal, parietal, and frontal cortices as the disease advances,^107^ with post-mortem studies revealing extensive neuronal and synaptic loss in cortical and subcortical regions that correlates with cognitive decline.^108^ This structural deterioration is paralleled by reduced glucose metabolism in temporoparietal areas, progressing to widespread cortical hypometabolism in later stages.^109–112^

The evidence reviewed so far highlights that AD progresses in two distinct phases: early-stage hyperexcitability and later-stage hypoactivity (Fig. 1b). M/EEG biomarkers have shown to be valuable tools for capturing these neural dysfunctions, offering a non-invasive window into the excitation-inhibition balance changes that underlie cognitive decline. However, while M/EEG provides crucial empirical insights, interpreting the causal mechanisms behind these alterations remains a challenge. Specifically, the precise contributions of Aβ and hp-*τ* pathology to neural hyper- /hypo-activity and subsequent network failure are still unknown. To address this gap, computational modeling approaches have been developed to integrate molecular, cellular, and network-level mechanisms of AD. These models allow us to simulate the neurophysiological consequences of AD-related pathology and test hypotheses about disease progression in a controlled and mechanistically interpretable manner. In particular, Neural Mass Models (NMMs) provide a powerful framework for understanding how AD-related disruptions in excitation-inhibition balance lead to oscillatory slowing, altered excitability, and ultimately, cognitive impairment.

NMMs are inspired by biological principles, and they offer a simplified yet robust framework for simulating the interactions of large populations of neurons. These models, which simulate the dynamics of excitatory and inhibitory neuron populations, provide insights into the impact of pathological conditions on neural activity and connectivity. For instance, studies such as those by de Haan et al. (2012),^113^ Alexandersen et al. (2023)^114^ and Sanchez-Rodriguez et al. (2024)^115^ have demonstrated how activity-dependent mechanisms contribute to hub vulnerability and oscillatory slowing in AD. Among the range of NMM-based approaches, many rely on established frameworks like Jansen-Rit^116^ or Wilson-Cowan models.^115^ While these models have advanced our understanding of single-frequency dynamics, particularly in the alpha range,^117, 118^ they are limited in addressing the multifrequency interactions critical to AD, even though some recent studies start incorporating theta-alpha interactions.^115^ The laminar neural mass model (LaNMM) proposed in Sanchez-Todo et al. (2023)^119, 120^ incorporates both alpha and gamma frequencies, which are intertwined and relevant for cognitive functions affected by AD. Unlike earlier models, it includes *PV* interneurons, and it also adds a physical layer that allows it to connect to physical recordings like EEG and MEG.

In this study, we extend the LaNMM to explicitly incorporate the core pathophysiological mechanisms of AD. By integrating both fast (gamma) and slow (alpha) oscillations within a single computational framework, the LaNMM captures the base dynamical oscillations that we need to explain the M/EEG biomarkers in AD progression. Specifically, we add to this model the implementation of Aβ oligomer-induced dysfunction of *PV* interneurons as a driver of early hyperexcitability and hp-*τ* -mediated hypoactivity as a key mechanism underlying later-stage network degradation. Unlike previous computational models that primarily focus on single-frequency alterations, the LaNMM accounts for the interplay between excitation-inhibition balance and oscillatory dynamics, together with a biophysical layer, enabling a direct comparison with empirical M/EEG biomarkers. Through this approach, we test the hypothesis that *PV* interneuron dysfunction alone is sufficient to produce the biphasic trajectory of AD-related spectral changes, but not the hypoactivity reflected in firing rate decreases, and that a synergistic interaction between Aβ toxicity and hp-*τ* pathology is required to fully capture disease progression (Fig. 1b). Ultimately, this work provides a mechanistically grounded computational framework for understanding the neural circuit dysfunction underlying AD and offers a valuable tool for testing hypotheses and guiding future experimental and clinical research.

## Materials and methods

### Laminar Neural Mass Model

The LaNMM (Laminar Neural Mass Model)^119, 120^ combines two well-established neural mass models, the Jansen-Rit (JR)^121^ and a modified Pyramidal Interneuron Gamma (PING) model^122, 123^ (Fig. 2a), to simulate slow and fast oscillatory dynamics within a cortical column. This hybrid approach integrates the strengths of each component: the JR model generates alpha-band (8–12 Hz) oscillations and operates at 10 Hz under baseline conditions, while the modified PING model produces gamma-band (30–70 Hz) oscillations with a baseline frequency of 40 Hz (Fig. 2c). The reciprocal coupling of these models allows the simulation of cross-frequency interactions, aligning with observed experimental data where alpha rhythms influence gamma activity.^120^ See a recent study which done a bifurcation analysis of the LaNMM where more frequency couplings (delta-gamma, theta-gamma, and alpha-gamma) are studied.^124^ The JR model consists of a population of Pyramidal neurons (*P*_1_), a population of excitatory interneurons (e.g., Spiny stellate cells *SS*), and a population of slow GABAergic interneurons (e.g., Somatostatin-expressing cells *SST*, such as Martinotti cells). The PING model consists of two populations, a Pyramidal (*P*_2_) and fast GABAergic interneuron (e.g., Parvalbumin-positive cells *PV*, such as basket cells). External inputs to the JR and PING models can represent inter-regional influences, allowing the model to capture realistic dynamics. The equations and parameters of the model are the ones in Sanchez-Todo et al. 2023.^120^ Following the mechanistic framework of reduced inhibition of fast interneurons (*PV*) on pyramidal cells, we represent the early progressive effects of the disease by reducing the connectivity from *PV* to *P*_2_. The hp-*τ* effects are modeled through the half of the maximum firing rate of the pyramidal populations 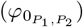. The model equations and parameters are the same as in Sanchez-Todo et al. (2023),^120^ and can be found in S1 Appendix.

**Fig 2.**
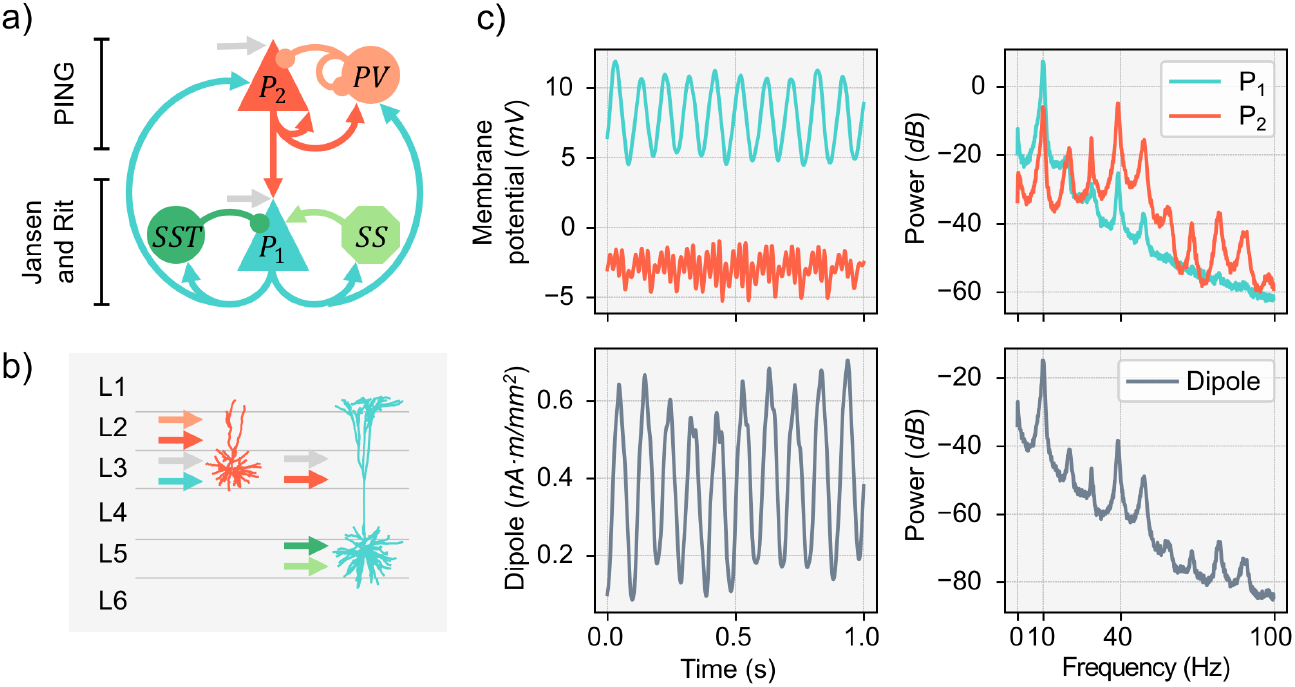
a) Illustration of the neuronal populations and the connectivity between them. Top, PING model, bottom, Jansen and Rit model. Rounded shapes represent inhibitory populations; the rest are excitatory ones. b) Biophysical Model. Geometries of the pyramidal cells used to generate compartmental models.^125^ The figure also shows the different depths at which the cortical layer divisions were set and the final synapse locations taken from Sanchez-Todo et al. (2023).^120^ c) Time series of the membrane potential of the pyramidal populations and the dipole, with the respective PSD profiles.

### Biophysical Model

To translate NMM activity into the equivalent dipole strength, we used a refined laminar mesoscale physical model,^125^ which was inspired by the laminar framework developed in Sanchez-Todo et al. (2023).^120^ In brief, a multi-compartmental modeling formalism and detailed geometrical models of pyramidal cells are used to simulate the transmembrane currents due to synaptic inputs into different cortical layers(Fig. 2b). These results are linearly combined to estimate the total current source densities (CSDs) generated due to the neural activity in pyramidal populations, which are considered the main current generators.^126, 127^ The CSDs are then used to calculate the equivalent dipole strength over time associated with the neural activity of the model.

### Spectral Features

The analysis of electrophysiological features in our study was conducted using time series of the dipole, simulated for a duration of 300 seconds, with a sampling frequency of 1000 samples per second. The numerical integration of the system was performed using the fourth-order Runge-Kutta method. To ensure robustness in our simulations using stochastic inputs, we employed 10 different noise seeds.

Spectral analysis was conducted to extract key frequency-domain features, including the alpha peak frequency, as well as power in the alpha (7–13 Hz) and gamma (30–60 Hz) bands. The estimation of these features was based on the power spectral density (PSD), computed using Welch’s method. The PSD was calculated over segments of length corresponding to ten times the sampling period and an overlap of five times the sampling period, following the established parameterization of Welch’s method. The peak frequency within the alpha band was identified as the frequency exhibiting the highest power within the 7–13 Hz range. Similarly, power in the alpha and gamma bands was computed as the summed spectral power density within the respective frequency intervals. The estimation of these features allows for a precise characterization of the oscillatory behavior of the modeled neuronal activity, facilitating a comparison with empirical electrophysiological findings.

### LaNMM implementation of pathological mechanisms

In this section, we outline how AD pathophysiology mechanisms can be implemented in the LaNMM and discuss the corresponding electrophysiological signatures at different AD stages. Moreover, we outline the rationale for the parameter changes that will be implemented to represent AD-related changes. Table 2 summarizes how each mechanism is mapped to the LaNMM, the parameter ranges used for the results, and the expected electrophysiological biomarkers according to the literature.

**Table 2.**
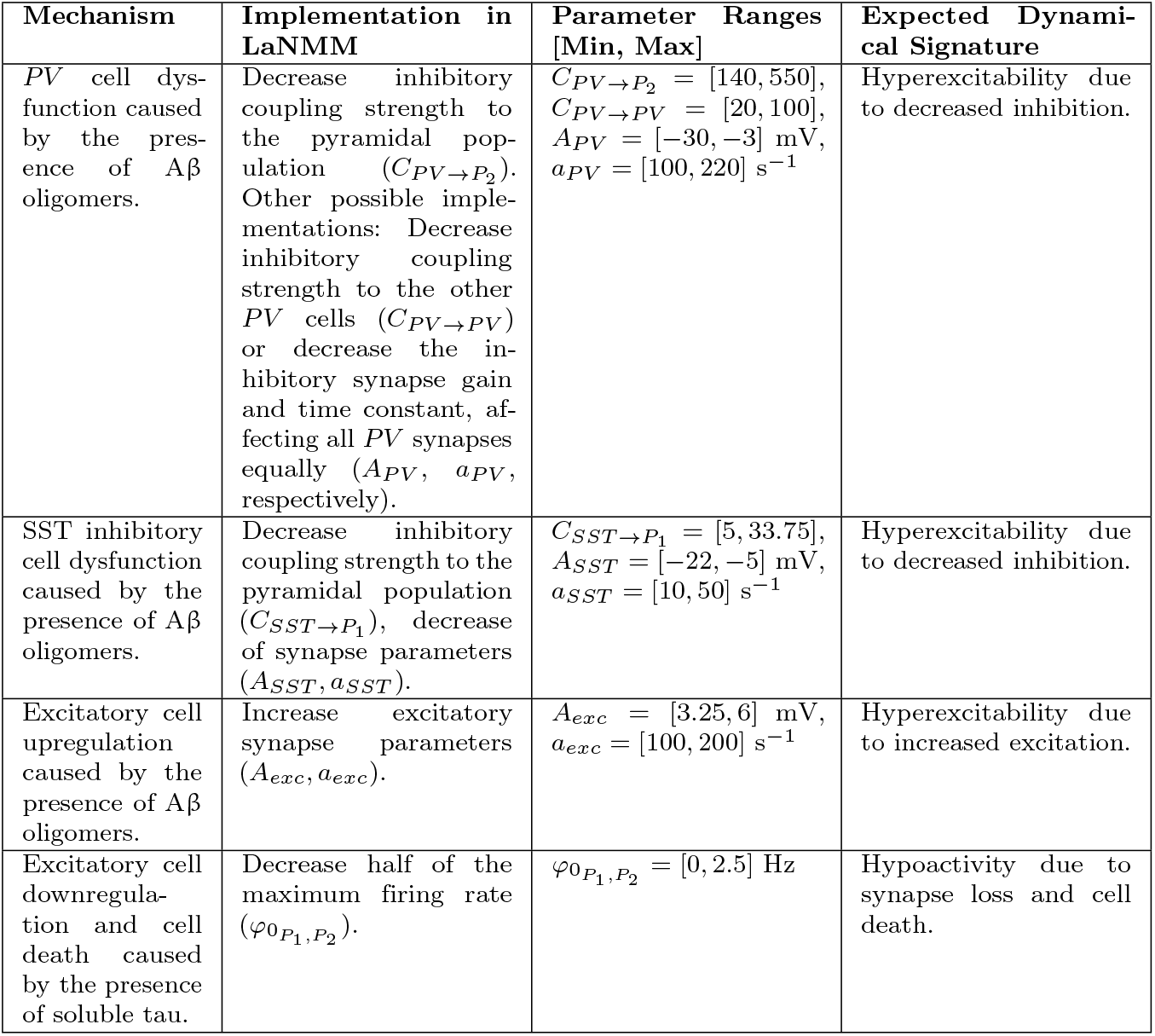
Mechanisms, model implementation, parameter ranges, expected dynamical signatures.

| Mechanism | Implementation in LaNMM | Parameter Ranges [Min, Max] | Expected Dynamical Signature |
| --- | --- | --- | --- |
| *PV* cell dysfunction caused by the presence of Aβ oligomers. | Decrease inhibitory coupling strength to the pyramidal population ($C_{PV \rightarrow P_2}$). Other possible implementations: Decrease inhibitory coupling strength to the other *PV* cells ($C_{PV \rightarrow PV}$) or decrease the inhibitory synapse gain and time constant, affecting all *PV* synapses equally ($A_{PV}$, $a_{PV}$, respectively). | $C_{PV \rightarrow P_2} = [140, 550]$,<br>$C_{PV \rightarrow PV} = [20, 100]$,<br>$A_{PV} = [-30, -3]$ mV,<br>$a_{PV} = [100, 220]$ s$^{-1}$ | Hyperexcitability due to decreased inhibition. |
| SST inhibitory cell dysfunction caused by the presence of Aβ oligomers. | Decrease inhibitory coupling strength to the pyramidal population ($C_{SST \rightarrow P_1}$), decrease of synapse parameters ($A_{SST}$, $a_{SST}$). | $C_{SST \rightarrow P_1} = [5, 33.75]$,<br>$A_{SST} = [-22, -5]$ mV,<br>$a_{SST} = [10, 50]$ s$^{-1}$ | Hyperexcitability due to decreased inhibition. |
| Excitatory cell upregulation caused by the presence of Aβ oligomers. | Increase excitatory synapse parameters ($A_{exc}$, $a_{exc}$). | $A_{exc} = [3.25, 6]$ mV,<br>$a_{exc} = [100, 200]$ s$^{-1}$ | Hyperexcitability due to increased excitation. |
| Excitatory cell downregulation and cell death caused by the presence of soluble tau. | Decrease half of the maximum firing rate ($\varphi_{0_{P_1, P_2}}$). | $\varphi_{0_{P_1, P_2}} = [0, 2.5]$ Hz | Hypoactivity due to synapse loss and cell death. |

To model the inhibitory dysfunction of *PV* neurons in the LaNMM, we gradually reduce the inhibitory coupling strength between *PV* and pyramidal cells, denoted as 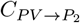. This reduction follows a cumulative distribution function, reflecting the progressive damage inflicted by Aβ oligomers (Fig. 3a). The presence of Aβ oligomers is represented by a Gaussian function, *G*(*x*), as suggested by Blomeke et al. (2024).^37^ Here, the variable *x* represents time and disease progression, ranging from a healthy state (S0, *x* = 0) to severe AD (*x* = 10). The mean of the Gaussian function is set at the transition between S2 and MCI (*µ* = 6), while Aβ oligomers begin accumulating at S1 (*x* = 2).^37^ The standard deviation is fixed at 1.5. The cumulative Gaussian function is then computed numerically via a discrete integral and normalized to range within [0, 1] (Fig. 3a).

**Fig 3.**
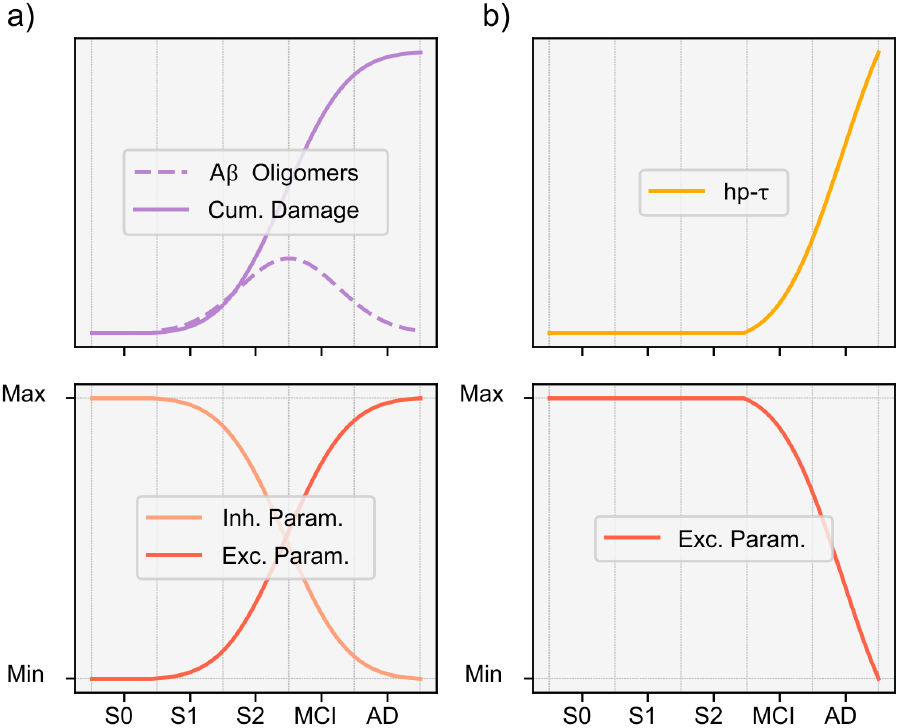
a) Top: Model of the Aβ oligomer concentration and its cumulative neurotoxic damage. Bottom: The variation of the model parameters is proportional to the cumulative damage. To simulate hyperexcitability resulting from reduced inhibition, inhibitory parameters are decreased from baseline. Conversely, to model hyperexcitability driven by increased excitation, excitatory parameters are upregulated in proportion to cumulative damage. b) Top: model of the hp-*τ* accumulation. Bottom: Excitatory parameters decrease in direct relation to hp-*τ* accumulation.

Separately, we conduct additional studies modifying other synaptic parameters, including the self-connectivity of *PV* cells (*C*_*P V* →*P V*_), the synaptic gain and time constant of *PV* interneurons (*A*_*P V*_, *a*_*P V*_) as well as SST interneuron parameters 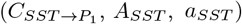. These adjustments enable a comparison with prior computational models of AD.^116–118^

To model hyperexcitability driven by increased excitation, we elevate the synaptic parameters (*A*_*exc*_, *a*_*exc*_) across all excitatory populations (*P*_1_, *P*_2_, *SS*). This adjustment follows the same cumulative distribution function to capture the cumulative effect of Aβ oligomers (Fig. 3a), reflecting the hyperexcitable state induced by Aβ in excitatory populations.

Conversely, to represent hypoactivity associated with hp-*τ*, we decrease the half of the maximum firing rate (*φ*_0_) of pyramidal populations (*P*_1_, *P*_2_). This decrease follows a sigmoid function peaking later in disease progression compared to Aβ accumulation, representing the delay between Aβ deposition and hp-*τ* pathology (Fig. 3b). This mechanism models neuronal loss, ultimately leading to an overall hypoactive state (a severe reduction of mean firing rate of the pyramidal cell population).

## Results

### *PV* dysfunction in LaNMM reproduces the oscillatory biomarkers of AD progression

To investigate the impact of Aβ oligomer accumulation on the model’s electrophysiological biomarkers, we systematically reduced the connectivity parameter from *PV* interneurons to *P*_2_ pyramidal cell populations 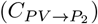, simulating the progressive interneuron dysfunction in AD (Fig. 4a). This reduction of inhibition not only induced an initial increase in oscillatory power but also reproduced later-stage loss of oscillatory activity and slowing of the alpha peak observed in AD.

**Fig 4.**
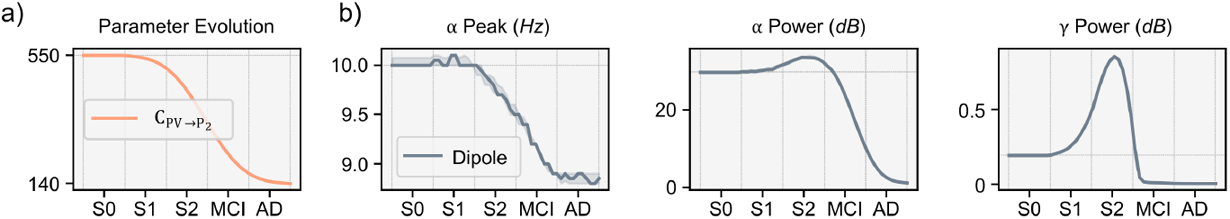
Electrophysiological biomarkers of the dipole model when altering the synapses from fast-spiking populations. a) The decrease of the connectivity parameter is proportional to the cumulative function of the Aβ oligomer concentration. b) Each column represents a different electrophysiological feature from the model’s dipole: *α* peak frequency, *α* band power and *γ* band power.

During the initial stages (S0, S1), oscillatory biomarkers (Fig. 4b) remained stable despite ongoing Aβ accumulation, which followed a cumulative function (Fig. 3a). As the oligomer load increased, a progressive decline in alpha peak frequency was observed, corresponding to the well-documented phenomenon of oscillatory slowing. Concurrently, alpha and gamma power initially increased, indicative of hyperexcitability due to the reduction in inhibitory control. Moreover, as the connectivity strength from *PV* interneurons to *P*_2_ populations continued to decline, a subsequent suppression of oscillatory activity emerged. This transition reflects the full continuum of AD-related spectral changes, progressing from an increase of spectral power to eventual oscillatory collapse. Our results reflect the critical role of *PV* inhibition failure in AD electrophysiology, aligning with prior empirical studies.

Thus, to fully capture the neurobiological complexity of advanced AD, the model should integrate additional mechanisms explicitly representing neuronal degeneration and loss. For that, we have computed the mean firing rate and membrane potential of the pyramidal populations and the generated dipole strength as the disease progresses, modeled by a reduction of 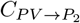 (Fig. 5). The results show that as the inhibitory connectivity decreases, there is an increase in the mean firing rate and the membrane potential, accompanied by a simultaneous decrease of the dipole mean strength. The increase in the firing rate might be expected in the initial stages,^81^ but not in the later ones^128^ when a generalized synapse loss is expected, and consequently, a decreased firing rate of the pyramidal populations (Fig. 1b).

**Fig 5.**
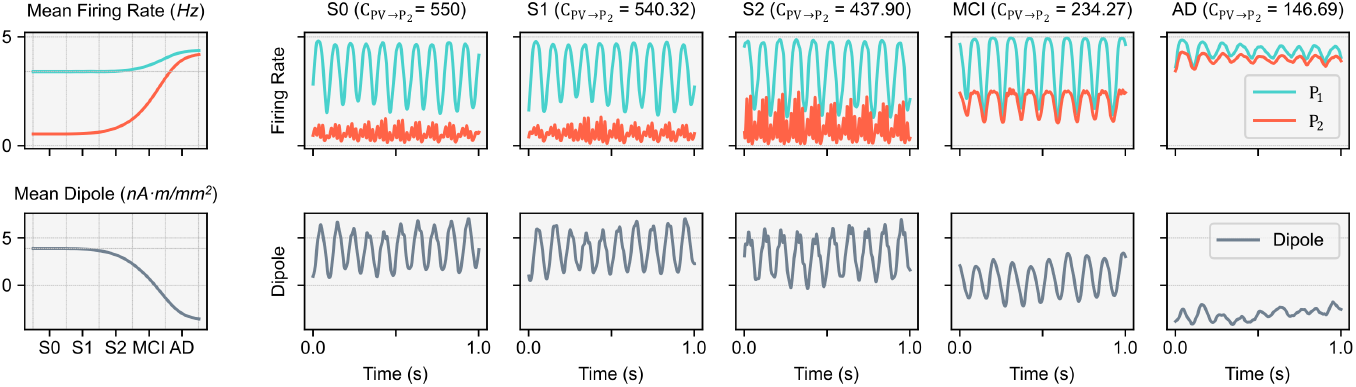
(left) Mean firing rate and dipole of the LaNMM as the 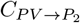 decreases, representing the cumulative damage of Aβ oligomers. (right) Samples of the time series of the firing rate and dipole for the different stages. Y-axis shared across rows.

For comparison with other parameters suggested in modeling studies and other parameters of the LaNMM, we also adjusted additional model parameters that might be influenced by Aβ oligomers and contribute to early-stage hyperexcitability (Fig. 6).

**Fig 6.**
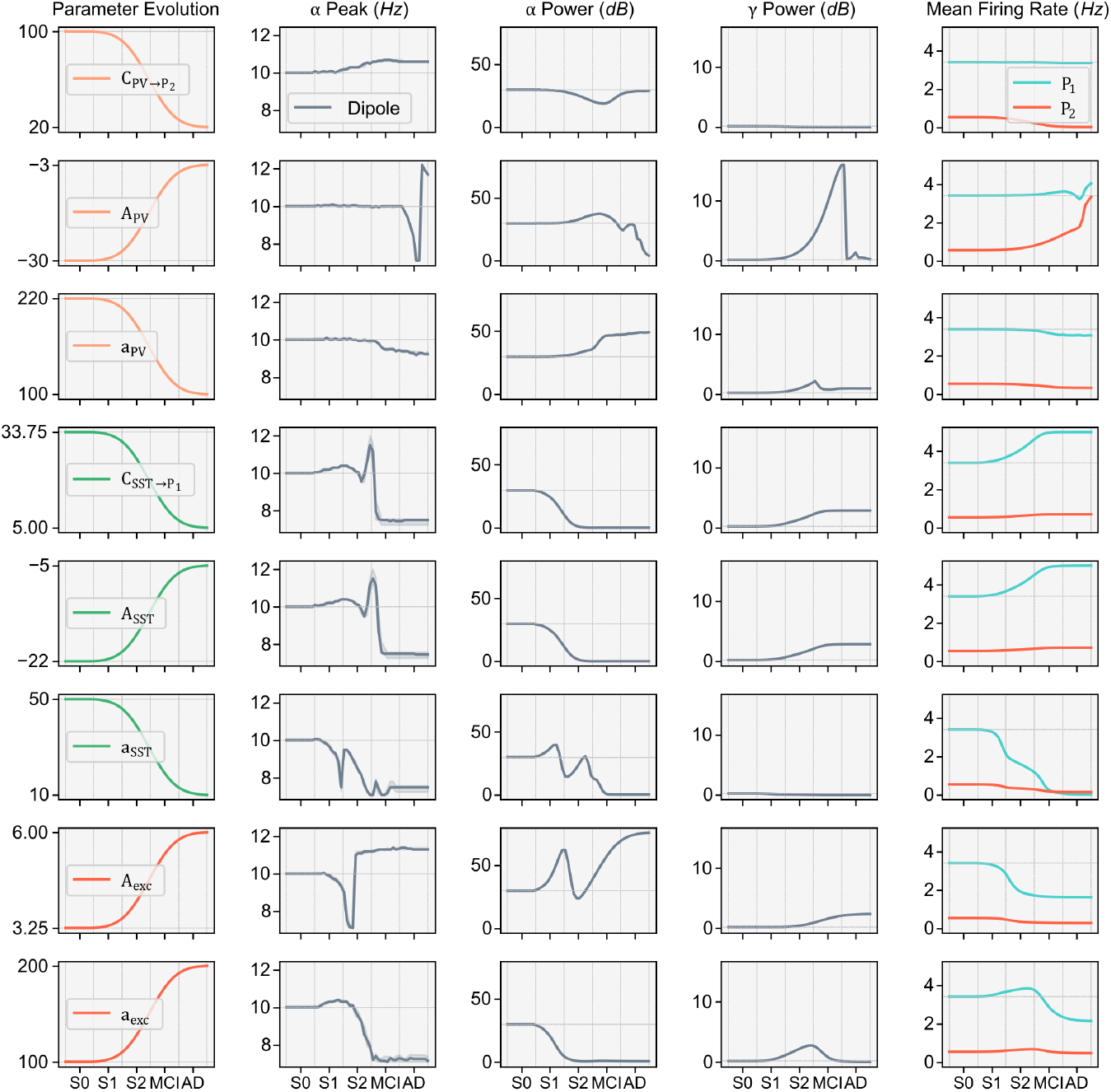
Electrophysiological biomarkers of the dipole model and the mean firing rates of *P*_1_ and *P*_2_ when altering different parameters that might be affected by the presence of Aβ oligomers (Fig. 3a). The first column shows the parameter changes, with their respective ranges Table 2. The first three rows represent a decreased inhibition of the *PV* cell population (color salmon), the next three of the *SST* inhibitory population (color green), and the last two rows represent increased excitation (color red).

To model other ways of *PV* cell damage by Aβ oligomers, we explored reductions in the self-connectivity of *PV* cells (*C*_*P V* →*P V*_) and reductions in the synapse parameters that affect all the *PV* synapses (*A*_*P V*_, *a*_*P V*_). None of the cases above replicated the biomarker progression as effectively as the connectivity from *PV* to 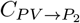, Fig. 4). Specifically, the synapse parameters *A*_*P V*_ and *a*_*P V*_ affect all the synapses of *PV* to other populations, as this parameters decrease, we can understand it as generalized cell damage or cell death, but none of the above seem to lead to the expected oscillatory biomarkers, only the 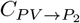 paramter.

Additionally, following previous modeling studies,^116, 117^ we modified the synaptic parameters of *SST* interneurons, but this also yielded unsatisfactory results (Fig. 6 and summary Fig. 9). Finally, we tested whether hyperexcitability could arise from increased excitation by adjusting all excitatory synapse parameters up, following the Aβ oligomer cumulative damage (*A*_*exc*_, *a*_*exc*_). However, none of these changes successfully reproduced the three-feature progression across AD stages (Fig. 6 and summary Fig. 9).

In summary, progressive reduction of inhibitory connectivity from *PV* interneurons to the pyramidal subpopulation *P*_2_ within the LaNMM reproduces the hallmark electrophysiological biomarkers of AD progression, including transient increases in *α* and *γ* power, slowing of the *α* peak frequency, and eventual suppression of oscillatory activity. This manipulation effectively captures the trajectory from early hyperexcitability to late-stage spectral collapse, underscoring the critical role of *PV* interneuron dysfunction in mediating the excitation-inhibition imbalance underlying AD network pathology. Notably, while this parameter captures the temporal evolution of spectral features(Fig. 1c), it does not lead to a reduction in the mean firing rate of pyramidal populations, which contradicts empirical observations of decreased neuronal activity in advanced AD(Fig. 1b). This discrepancy highlights the necessity of incorporating additional mechanisms—such as synaptic loss or reduced intrinsic excitability—to more fully account for the neuronal hypoactivity characteristic of later disease stages. Finally, among the various parameter perturbations tested, only the reduction of 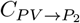 successfully recapitulated the full spectrum of electrophysiological alterations, suggesting a selective vulnerability of this inhibitory pathway in AD-related network dysfunction.

### Combined model of interneuron and pyramidal cell dysfunction explains electrophysiological dysregulation across disease progression

Our current model of *PV* interneuron dysfunction accurately reproduces key electrophysiological biomarkers observed during AD progression, particularly changes in oscillatory power and peak frequency (Fig. 4 mapped to Fig. 1c). However, it does not inherently account for structural findings such as neuronal loss documented by MRI or postmortem studies(Fig. 5 mapped to Fig. 1b). Specifically, the model predicts sustained neuronal hyperactivity at advanced stages (modeled by a reduction of 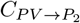, with increasing firing rate in the pyramidal populations, which is inconsistent with actual cell loss (Fig. 5).

We have developed a model of the combined effects of Aβ oligomers and hp-*τ* by altering the 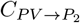 parameter as in Fig. 4 (Fig. 7 top row), and modifying the half of the maximum firing rate constant of the pyramidal populations, *φ*_0_ (Fig. 7 middle row), to represent the hp-*τ* pathology associated pyramidal cell loss, following the distribution in Fig. 3b. By varying only the *φ*_0_ parameter, following the hp-*τ* damage across stages, it is only possible to reproduce the biphasic progression in the gamma power band but not in the rest of the features (alpha power and peak). By combining both parameters (Fig. 7 bottom row), the model does not only reproduce the M/EEG biomarker signatures across disease progression but also the initial increase and then decrease of FR (Fig. 5 mapped to Fig. 1b) associated with cell loss. See Fig. 9 for a summary fo the mechanisms implemented, the specific parameter changed in the LaNMM to represent the mechanisms, and the success or failure to reproduce the specific electrophysiological features of AD progression across stages.

**Fig 7.**
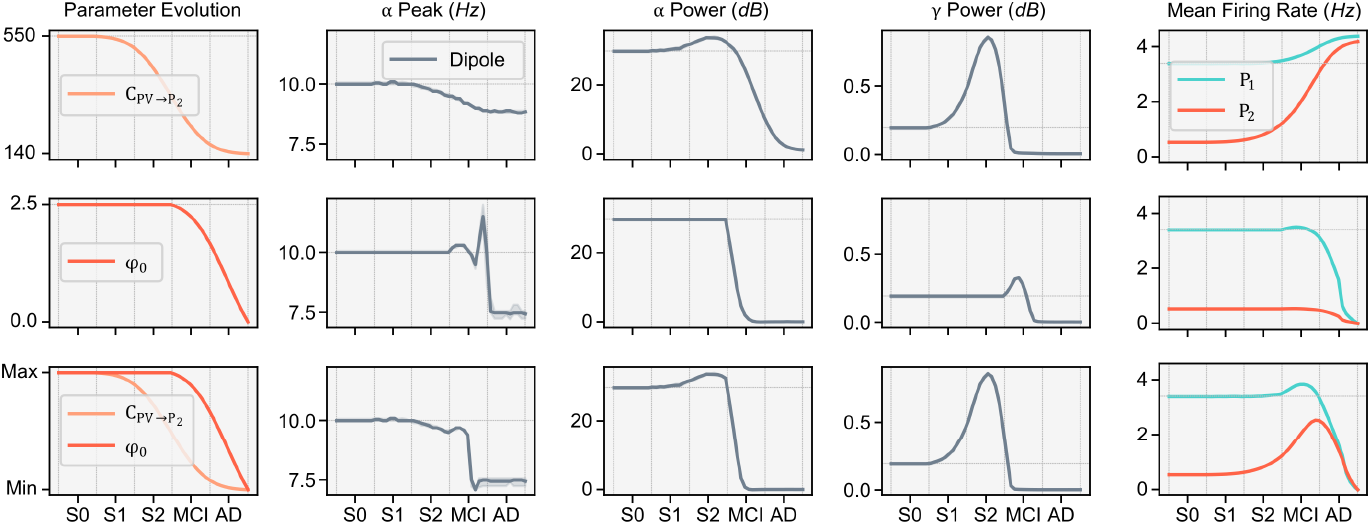
Electrophysiological biomarkers of the dipole model and mean firing rates of *P*_1_ and *P*_2_ when altering the connectivity parameter from the *PV* to *P*_2_ cells 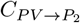, top row), the half of the maximum firing rate of the pyramidal cells (*φ*_0_ of *P*_1_ and *P*_2_, middle row), and the combination of both (bottom row).

**Fig 8.**
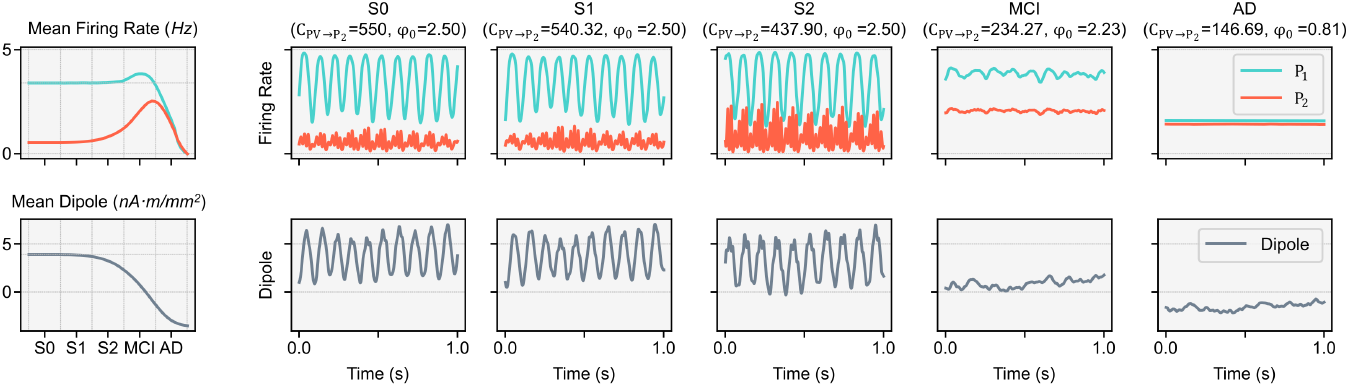
(left) Mean firing rate and dipole of the LaNMM as the 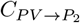 and *φ*_0_ decreases representing the cumulative damage of Aβ oligomers and hp-*τ* respectively. (right) Samples of the time series of the firing rate and dipole for the different stages. Y-axis shared across rows.

**Fig 9.**
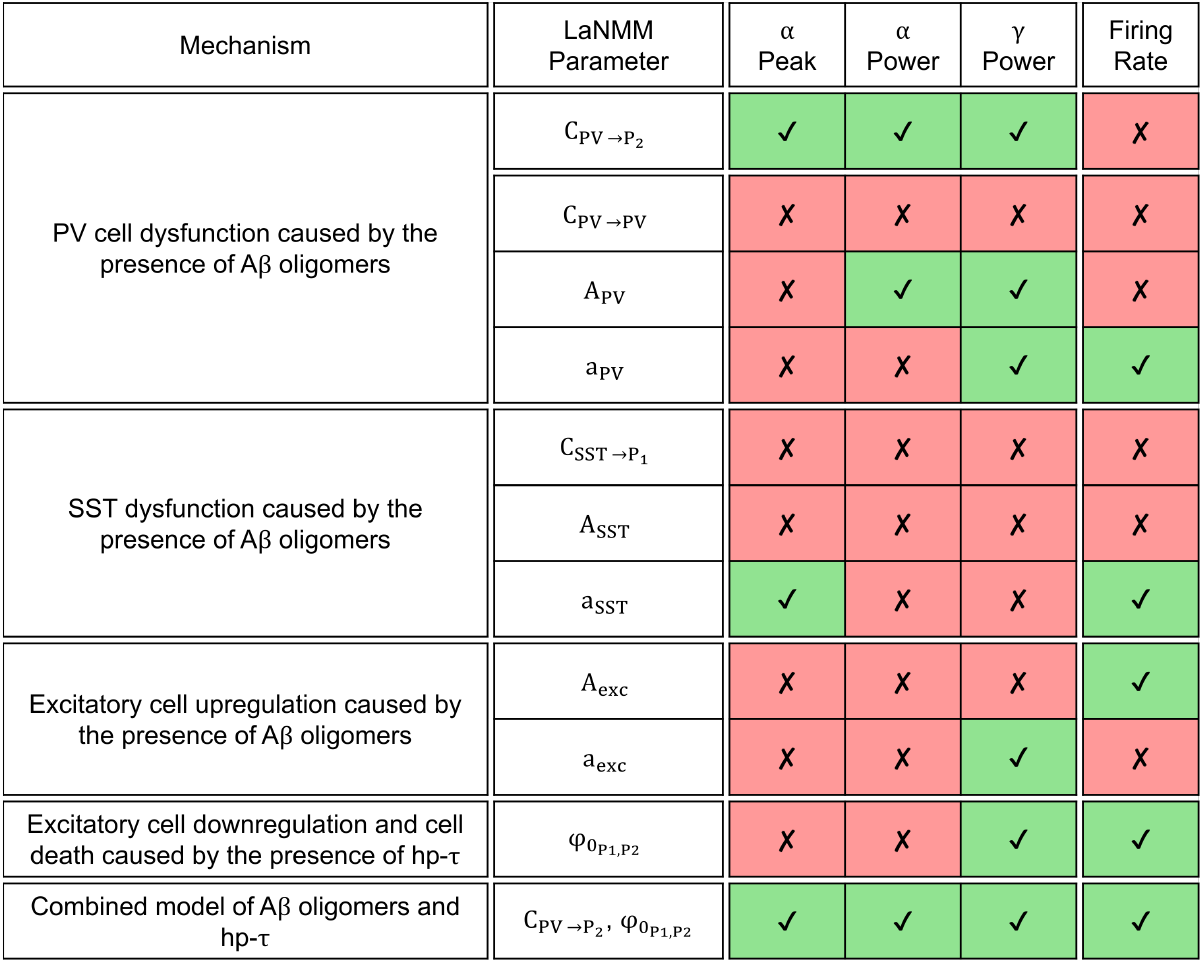
Summary table of the mechanisms implemented in the LaNMM, the specific parameter altered, and the success (✓) or failure (✗) in representing the electrophysiological biomarkers of AD across the disease progression.

These findings suggest that *PV* interneuron dysfunction is a primary driver of early-stage hyperexcitability, whereas hp-*τ* -induced neuronal loss (modeled with a decrease of 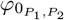) underlies the later-stage transition to hypoactivity (Fig. 1b). The observed biphasic progression, characterized by an initial rise and subsequent decline in alpha and gamma power and the slowing of the alpha peak, aligns with empirical EEG/MEG data, strengthening the role of computational modeling in elucidating AD pathophysiology.

## Discussion

Our findings demonstrate that the progression of AD can be effectively modeled using the LaNMM,^119, 120^ by integrating key pathophysiological mechanisms of the disease through parameter adjustments. A distinct advantage of this model is its combination of both fast (PING) and slow (Jansen-Rit) circuits, enabling the representation of critical neural players in AD, particularly *PV* interneurons. Furthermore, because the laminar model can be used to simulate dipole activity through a biophysical framework,^125^ it allows for direct comparison with empirical EEG and MEG data. By systematically adjusting the inhibitory connectivity of fast-spiking *PV* interneurons and the mean firing rate of pyramidal cells, we successfully replicated, in a physiologically meaningful way, the biphasic electrophysiological trajectory of AD—an initial phase of hyperexcitability followed by oscillatory slowing and hypoactivity.

The results align with prior neurophysiological evidence indicating that Aβ oligomers induce early-stage cortical hyperactivity by impairing *PV* interneuron function.^45, 47, 48, 81–84^ This dysfunction leads to increased gamma and alpha power, mirroring empirical M/EEG findings in preclinical AD.^4, 60^ The model further shows how, by representing *PV* cell dysfunction associated with a Aβ oligomer toxicity, inhibitory control diminishes, leading to electrophysiological changes along the trajectory of AD consistent with findings from EEG studies in MCI and AD.^61, 72^

We examined alternative mechanisms for hyperexcitability, including increased excitatory drive,^38, 88^ but found that *PV* interneuron dysfunction best explained the observed oscillatory signatures. Notably, when modeling Aβ oligomer-induced damage to *PV* interneurons alone, we observed an initial hyperexcitability phase. This increase in excitation is a direct consequence of desinhibition in the model.

To produce the expected progressive reduction in firing rate seen in later disease stages associated with hp-*τ*, we represented pyramidal cell hypoactivity via its mean firing rate setting parameter 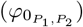, which simply captured the progressive decline in neural activity. This result supports the notion that early-stage AD is predominantly driven by an imbalance between excitation and inhibition, with a specific vulnerability of *PV* interneurons.^45, 47, 83^ Furthermore, by incorporating hp-*τ* -induced hypoactivity in later stages,^99, 100^ our model successfully captures the transition from hyperexcitability to hypoactivity and cell loss, paralleling clinical progression.

A critical implication of our findings is that *PV* interneuron dysfunction may serve as an early biomarker for AD, detectable through M/EEG before significant hp-*τ* deposition occurs. This aligns with studies demonstrating that gamma-band abnormalities precede cognitive decline and correlate with Aβ pathology.^48, 84^ Additionally, our results reinforce the importance of preserving inhibitory function as a potential therapeutic target,^83^ including sensory^129–131^ or transcranial electrical stimulation.^132–134^ Recently, the LaNMM has been used in a whole-brain model to model how oscillatory dynamics in AD have the potential to be restored using psychedelics.^135^

Despite these insights, limitations remain. Our model does not account for regional variations in pathology, nor does it fully integrate vascular and glial contributions to network dysfunction. Future work should incorporate multimodal data into a whole-brain model based on the LaNMM to refine mechanistic interpretations and validate model predictions against longitudinal patient cohorts.

Finally, while our work has focused on AD, our model of *PV* dysfunction is relevant to other conditions, such as ASD and Schizophrenia. Interneuron dysfunction, particularly involving *PV* interneurons, is implicated in autism spectrum disorder (ASD) and Schizophrenia.^136^ In schizophrenia, reduced *PV* interneuron function leads to disrupted gamma oscillations, impairing sensory processing and predictive coding.^137^ Similarly, in ASD, alterations in *PV* interneurons contribute to excitatory/inhibitory imbalances, affecting neural oscillations and predictive coding.^137^ These findings suggest that *PV* interneuron dysfunction across these disorders compromises neural oscillations and leads to cognitive deficits.

## Conclusion

In summary, our study demonstrates that the LaNMM can effectively reproduce the biphasic electrophysiological trajectory of AD, encompassing an early phase of hyper-excitability due to Aβ-induced parvalbumin-positive (*PV*) interneuron dysfunction, followed by oscillatory slowing and hypoactivity driven by hyperphosphorylated hp-*τ* -related pyramidal cell hypoactivity. This modeling framework successfully bridges molecular pathology and large-scale neural dynamics, replicating key M/EEG biomarkers and providing mechanistic insight into their emergence through targeted alterations in inhibitory connectivity and excitatory drive. The implications of our findings extend beyond theoretical modeling; they reinforce the central role of *PV* interneurons in early-stage AD, highlighting them as promising therapeutic targets for interventions aimed at restoring inhibitory control. Moreover, the integration of Aβ and hp-*τ* mechanisms within a unified model underscores that network degradation results not from isolated pathological events but from their dynamic interaction across cortical circuits. Future directions include incorporating patient-specific data into whole-brain instantiations of the LaNMM to account for individual variability in Aβ and hp-*τ* burden, as well as extending the model to encompass additional pathophysiological contributors such as neuroinflammation and vascular dysfunction. The framework also offers a platform for simulating potential therapeutic strategies—pharmacological or neurostimulation-based—to evaluate their predicted effects on disease progression and inform precision intervention approaches.

## Supporting information

### S1 Appendix. LaNMM model equations and parametes

Neural mass models (NMM) mathematically represent the dynamics of the average membrane potential and firing rate of a neuronal population in a cortical column.^138^ These models are described by a second-order differential equation that defines the average membrane perturbation *u*_*m*←*n*_(*t*) a population *m* experiences at synapses from another population *n*. The relationship between the presynaptic firing rate *φ*_*n*_ and the postsynaptic potential is governed by the operator 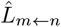 and its inverse:

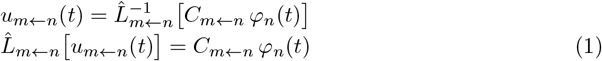

Here, *C*_*m*←*n*_ is the connectivity constant, and 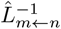 is expressed as a convolution with a kernel *h*(*t*) = *Aat* exp[−*at*] for *t >* 0, satisfying the Green’s function equation. ^139^ For generality, synapse *s* (*m* ← *n*) dynamics are defined as:

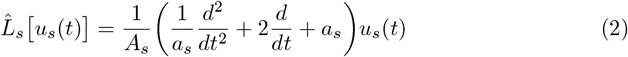

where *A*_*s*_ is the synaptic gain, and *a*_*s*_ = 1*/τ*_*s*_ is the synaptic rate constant. The summed perturbations *v*_*m*_ from incoming synapses produce a nonlinear firing rate *φ*_*m*_ via a sigmoid function:

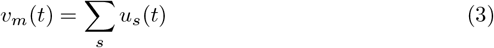

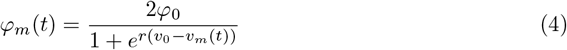

Here, *φ*_0_ is half the maximum firing rate, *v*_0_ is the potential at *φ*_0_, and *r* determines the sigmoid’s slope.

The parameters of the model are described in Table 3, and the equations are the following:

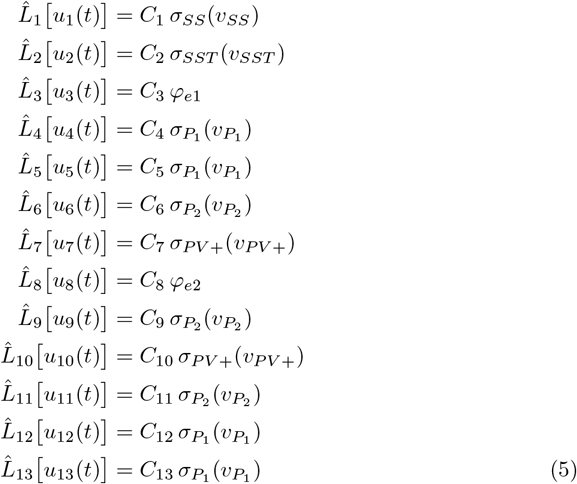

with neuronal population membrane potentials given by:

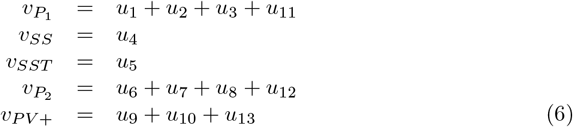

**Table 3.**
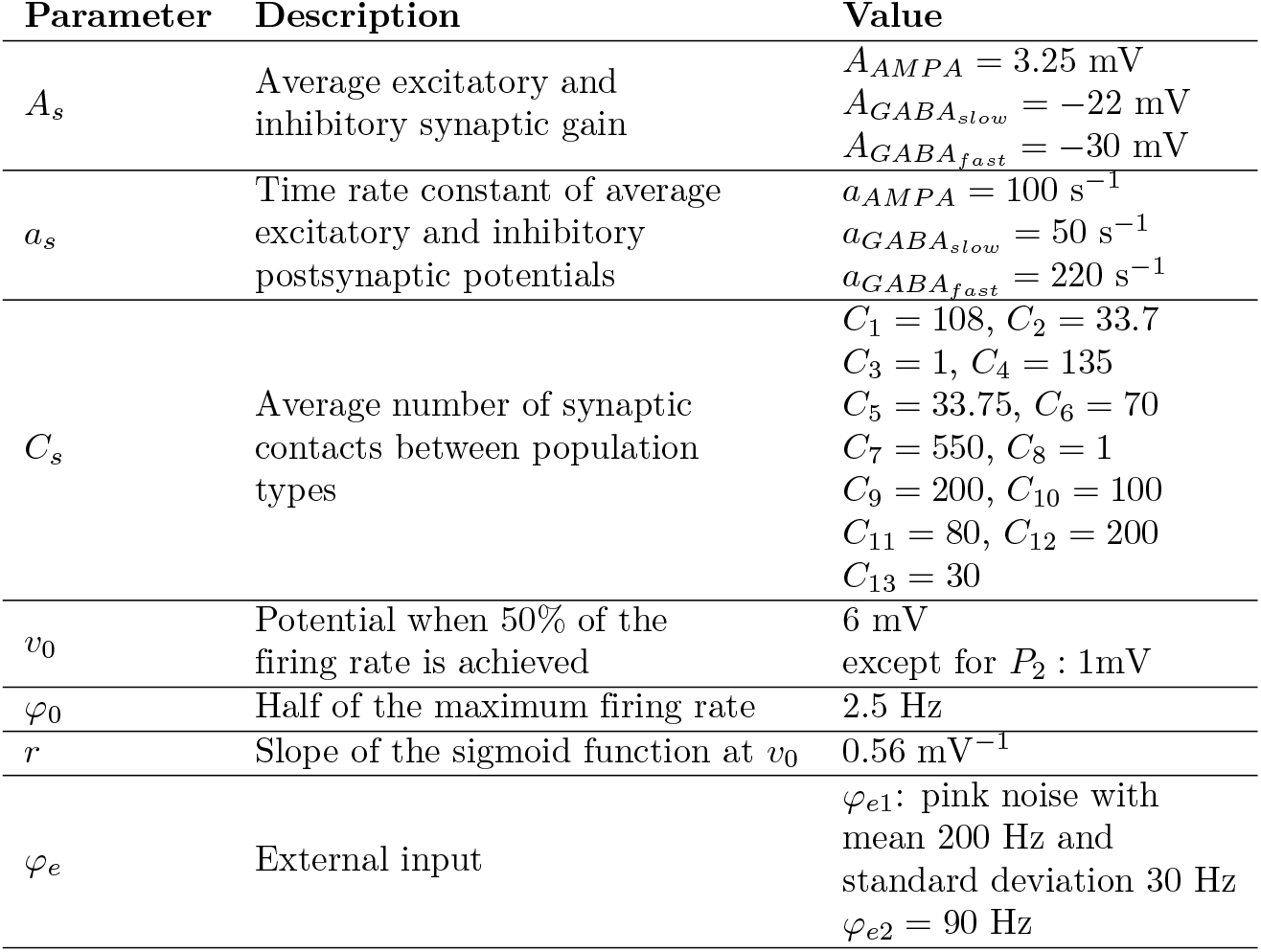
Parameters, description, and standard values of the model. Values are taken from.^120^ Note that the sigmoid parameters in this model (*v*_0_, *φ*_0_, *r*) are common to all neuron populations. Moreover, excitatory synapses have the same synapse dynamics, 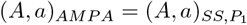, and inhibitory synapses have either fast dynamics, 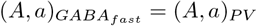, or slow, 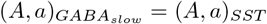.

## Acknowledgments

Roser Sanchez-Todo, Borja Mercadal, Edmundo Lopez-Sola, Maria Guasch-Morgades, Gustavo Deco, and Giulio Ruffini are funded by the European Commission under European Union’s Horizon 2020 research and innovation programme Grant Number 101017716 (Neurotwin) and European Research Council (ERC Synergy Galvani) under the European Union’s Horizon 2020 research and innovation program Grant Number 855109.

Special thanks to Elia Lleal, Raul de Palma Aristides, Jakub Vohryzek, and Adri`a Galan-Gadea for the revision of the text and additional support.

